# Placebo treatment entails resource-dependent downregulation of negative inputs

**DOI:** 10.1101/2023.09.05.556222

**Authors:** Jonas Rauh, Arasch Mostauli, Matthias Gamer, Christian Büchel, Winfried Rief, Stefanie Brassen

## Abstract

**Background:** Clinical trials of antidepressants show improvements in placebo groups of up to 80% compared to the real treatment arm. The mechanism underlying this clinically important effect has been linked to expectation induced goal-directed control. Here, we investigated how cognitive resources influence the effects of positive expectations on emotional processing.

**Methods:** Forty-nine healthy volunteers participated in a cross-over fMRI study, in which expectancy of positive emotional changes was induced by an alleged oxytocin nasal spray combined with verbal instruction. Participants performed a spatial cueing paradigm that manipulated the attention to emotional face distractors in the scanner and were characterized regarding their general ability to control attention.

**Results:** Behavioral findings showed placebo treatment to improve mood and to reduce distractibility by fearful over happy faces, specifically when more attentional resources were available to process faces. This aligned with neural changes in activation and functional coupling in lateral prefrontal-limbic networks indicating an expectation induced top-down regulation of aversive inputs. In addition, behavioral effects and prefrontal-parietal engagement directly correlated with trait ability to control attention.

**Conclusions:** Our data show that placebo treatment combined with verbal instruction alone can improve mood and recruit top-down attentional selection. Changes in emotional processing critically depended on attentional context and individual control ability (i.e., contextual and general resources). These findings may be particularly relevant in patients with major depressive disorder, who often demonstrate a negativity bias and in whom placebo effects by verbal instructions alone may be limited due to reduced cognitive control capacity.

## Introduction

There is an ongoing debate about the influence of patient expectations on the effectiveness of antidepressant treatment, prompted by studies showing that placebo groups can achieve up to 80% of the benefits observed in groups receiving antidepressants (1–6). Insights into the mechanisms of placebo effects on mood state and emotion processing have the potential to improve treatment strategies for depression. However, identifying mechanisms underlying placebo responses requires contrasting the placebo condition with a non-expectation control condition.

There is solid evidence that placebo treatments can substantially modulate affect (7,8). For instance, positive expectations can reduce experimentally induced sadness (9–11), anxiety or fear (12–16) and social stress (17,18). Moreover, placebo manipulations can enhance baseline mood states (16,19). Interestingly, effects persist in open-label placebo studies (20–22).

Imaging studies of placebo effects in the affective domain often report an involvement of the lateral prefrontal cortex (PFC) (14,23), the ventromedial PFC (vmPFC) (24), and the anterior cingulate cortex (ACC) (11,14,16). Prefrontal engagement has been attributed to the construction and maintenance of instructed beliefs (25,26). In particular, the lateral PFC seems to be pivotal for building and updating of treatment expectations, and the signaling of instructed state representations to bias perceptual processing in downstream networks (26,27).

The consistent finding of prefrontal engagement in expectation effects suggests a critical role of cognitive functioning and goal-directed control (28). However research on the impact of cognitive function on placebo effects is heterogeneous (29–31) and mainly focused on the pain system. Far less is known about cognitive contributions to affective expectation effects, despite its relevance to clinical conditions like major depression, where prefrontal and executive functions are frequently compromised (32–35).

Here, we combined the induction of positive expectations with functional neuroimaging of prefrontal networks during emotional processing. Forty-nine healthy volunteers, characterized in their general cognitive control ability, participated in a randomized, cross-over fMRI study where they performed a spatial cueing paradigm that manipulated attentional resources available to process emotional face distractors in a goal directed manner (36,37). Thus, cognitive resources were operationalized as both state/contextual and general variables.

We hypothesized that positive expectations improve mood state and bias emotional processing of faces. We expect these effects to depend on contextual and general resources of cognitive control and to be modulated by prefrontal-limbic networks.

## Methods and Materials

### Participants

Fifty-five volunteers were recruited for the study, all free from psychiatric or neurological disorders, and medication. Technical issues led to the exclusion of two participants, three did not believe in receiving oxytocin from the beginning, and one exhibited strong movement artifacts, resulting in a final sample size of N = 49 (mean age: 26.92 ± 4.23 years; 30 women). The local ethics committee approved the study (Ethikkommission der Ärztekammer Hamburg). All participants gave written informed consent and were financially compensated for participation. The study has been registered at the German Clinical Trials Register (https://drks.de/search/en/trial/DRKS00027289).

### Experimental protocol and expectation induction

Participants attended three study days (Figure 1A). On day 0, they underwent screening and completed a visual search task (Singleton task, see below). On day 1 and 2, separated by 1 week, participants underwent fMRI scanning. Before scanning, they received a saline nasal spray, labeled either as oxytocin (condition ‘placebo’) or saline (condition ‘control’) in a counterbalanced cross-over design (Figure 1A). Expectations about positive oxytocin effects (e.g., “Oxytocin can improve mood and inhibit the perception of aversive stimuli such as fearful faces”) were induced by a custom 5-minute video documentary. Participants watched the video on both days before being told whether they were in the placebo or control condition that day. Participants then self-administered a saline nasal spray, dispersing four puffs, two per nostril.

**Figure 1.**
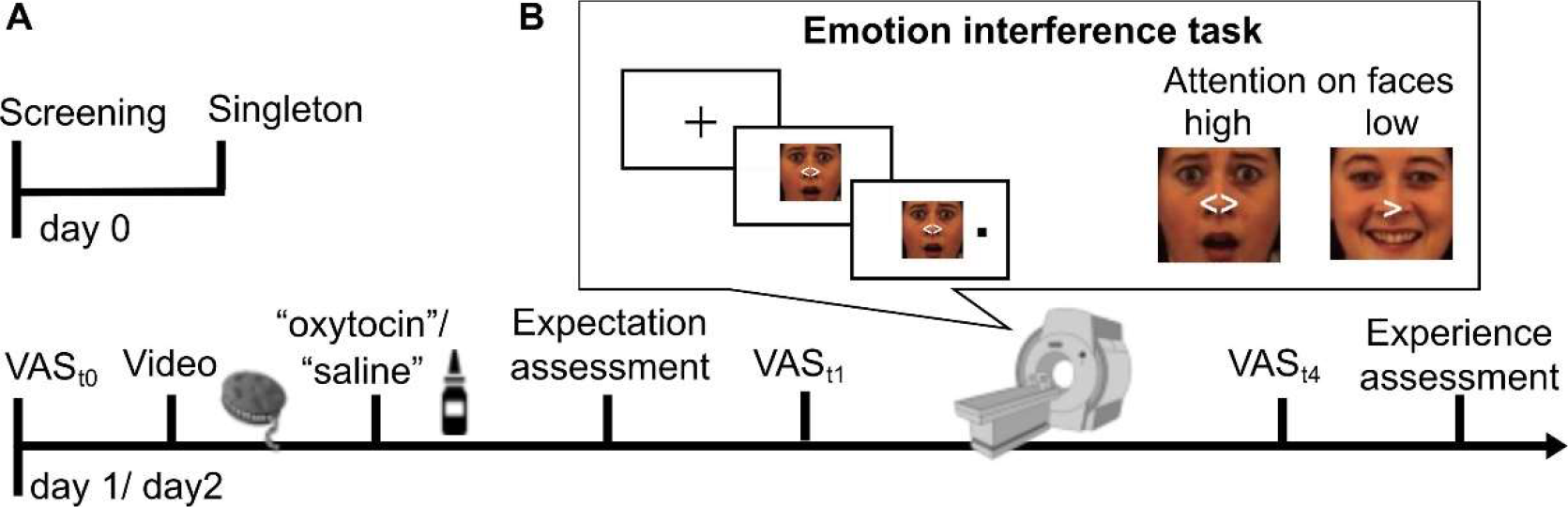
Study design and paradigm. **(A)** Timeline. **(B)** Emotion interference task with example stimuli depicting trials with high (non-directed cue) and low (directed cue) attentional resources for processing face distractors. VAS = visual analogue scale.

### Behavioral ratings

Mood state was assessed using a visual analogue scale (VAS) at participants’ arrival (baseline VAS_t0_), after nasal spray application, and after each of the three scanning blocks (VAS_t1-4_). Participants had to shift a randomly positioned cursor on a scale spanning from ‘unhappy’ to ‘happy’ with 400 incremental steps. Values were baseline corrected by subtracting VAS_t0_ from VAS_t1-4_ values.

Expectation of positive mood change was assessed after nasal spray application using an 11-point-scale (0 for no expected mood enhancement and 10 for large expected mood enhancement). At the end of each study day, participants used the same scale (0 for no and 10 for a significant positive mood improvement) to indicate how they had actually experienced the mood change (both scales adapted from (38)).

### Emotional interference task

In the scanner, participants performed a modified spatial cueing paradigm established in our laboratory (36,37). Participants had to respond to left- or right-sided dot-targets, preceded by spatially informative or uninformative cues (Figure 1B). After a 1000 ms fixation, cues appeared on neutral, happy, sad, fearful and scrambled face distractors (5° of visual angle) for 2050-2500 ms. The left- or right-sided dot target (horizontally displaced by 7°) followed for 200 ms. Participants were instructed to respond quickly and accurately by button press according to target side. They were told to ignore the faces and instructed to keep fixation throughout the task. Trials ended with a randomly chosen inter-trial-interval of 2000-3500 ms. In two-thirds of trials, a white arrowhead pointed to the target side (valid, 83%) or the opposite side (invalid, 17%). The few invalid trials aimed to maintain attention and were excluded from analyses. In one-third of all trials, a neutral, non-spatial cue indicated equal target likelihood on both sides. As demonstrated previously, without a directional cue, the attentional focus is not covertly shifted, allowing more resources for controlled face processing according to emotional preferences (36,37).

Three sessions comprised a total of 360 trials, with 1-minute breaks in between. The fully counterbalanced design had a pseudo-randomized trial order. The stimuli included 18 facial identities (9 female) from the well-validated Karolinska Directed Emotional Faces set (39), each displaying four emotional expressions, and scrambled pictures created by shuffling the luminance of the neutral face. Each picture was presented four times. Participants were trained on the task before scanning. After scanning, they rated valence and arousal (range: 1-4) for all presented faces to account for potential differences in valence categorization and arousal perception between expressions and sessions. Behavioral tasks utilized the Psychophysics Toolbox (PTB, 3.0.14) for MATLAB.

### Eye movement recording

Central fixation was instructed throughout the experiment and controlled by an online infrared eye-tracker (Eyelink 1000, SR Research), oriented toward the right eye. Gaze positions were sampled at 1000 Hz, allowing accurate saccade and fixation tracking. Stimulus onsets were recorded in parallel to the eye-tracking data. For details on the analysis see Supplementary information.

### Singleton task

On the screening day, participants performed a visual search task on a computer that measures participants’ ability to recruit top-down control in order to flexibly focus on or to ignore salient visual stimuli (“singleton detection mode”) (40–42). Performance in this task has previously been related to individuals’ ability to allocate attentional resources trial-by-trial in the service of prioritized emotional goals (41). For details of the task and the calculation of the Singleton score see Supplementary information.

### MR image acquisition

Imaging data were acquired on a Siemens PRISMA 3T scanner (Erlangen, Germany) using a 64-channel head coil. Multiband echo-planar imaging sequences were applied to obtain the functional imaging data (66 slices, voxel size 1.5 × 1.5 × 1.5 mm, 1.55 s TR, 29 ms TE, 70° flip angle, 225 mm field of view, multiband mode with 3 bands). A T_1_-weighted MPRAGE anatomical image was also obtained for functional preprocessing (voxel size of 1 × 1 × 1 mm, 240 slices).

### Image preprocessing and analysis

Image preprocessing and analysis were performed utilizing SPM12 (Wellcome Trust Centre for Neuroimaging, London, UK) and custom scripts in MATLAB. Preprocessing included realign and unwarp, normalization utilizing T1 structural image information, and 4-mm smoothing. The TsDiffAna toolbox (http://www.fil.ion.ucl.ac.uk/spm/ext/#TSDiffAna) was used for image diagnostics.

Statistical analysis applied a two-level random effects approach employing the general linear model. At single subject level, onsets of all 10 conditions (5 emotions × 2 cues) and of invalid and incorrect trials (regressor of no interest) were modeled as separate regressors convolving delta functions with a canonical hemodynamic response function. Data from the placebo and control condition were defined as separate sessions and entered into a single model. To enhance signal-to-noise ratio, individual noise regressors derived from the GLMdenoise toolbox (https://kendrickkay.net/GLMdenoise/, Version 1.4), were entered as regressors of no interest into the first-level model.

Individual contrast images were then entered into second-level random-effect ANOVA models including the factors stimulus (all emotions for confirmatory analyses, happy and fearful faces for target analyses), cue and session. A simple regression model was applied for correlation analysis.

Functional connectivity was assessed using a generalized psychophysiological interaction analysis (gPPI toolbox; http://www.nitrc.org/projects/gppi). The time course from 2mm spheres was extracted around individual peaks within the identified cluster. On the first level, adjusted time series were modeled together with onset vectors and the interaction of both. Specific interaction contrast images (i.e., fearful/happy, non-directed/directed, placebo/control) were then analyzed within second level factorial designs.

We report results corrected for FWE due to multiple comparisons. Correction was conducted at the peak level within small volume ROIs for which we had an a-priori hypothesis or when passing a whole-brain corrected cluster threshold of FWE < .05 (cluster forming threshold p < .001 uncorrected).

Based on our main interest in prefrontal-limbic regulation (14,26), we created ROIs of the dlPFC, the vmPFC/ACC, and the amygdala using functional clusters from meta-analyses on neurosynth.org and anatomical masks from the AAL3 atlas (43) (Supplementary Figure 1).

## Results

### Positive expectation induction enhances mood state and post-treatment valuation

Participants rated their expectation of positive mood effects as significantly higher in the placebo than in the control session (T_(48)_ = 7.90, p < .001, d = 1.13, Figure 2A).

**Figure 2.**
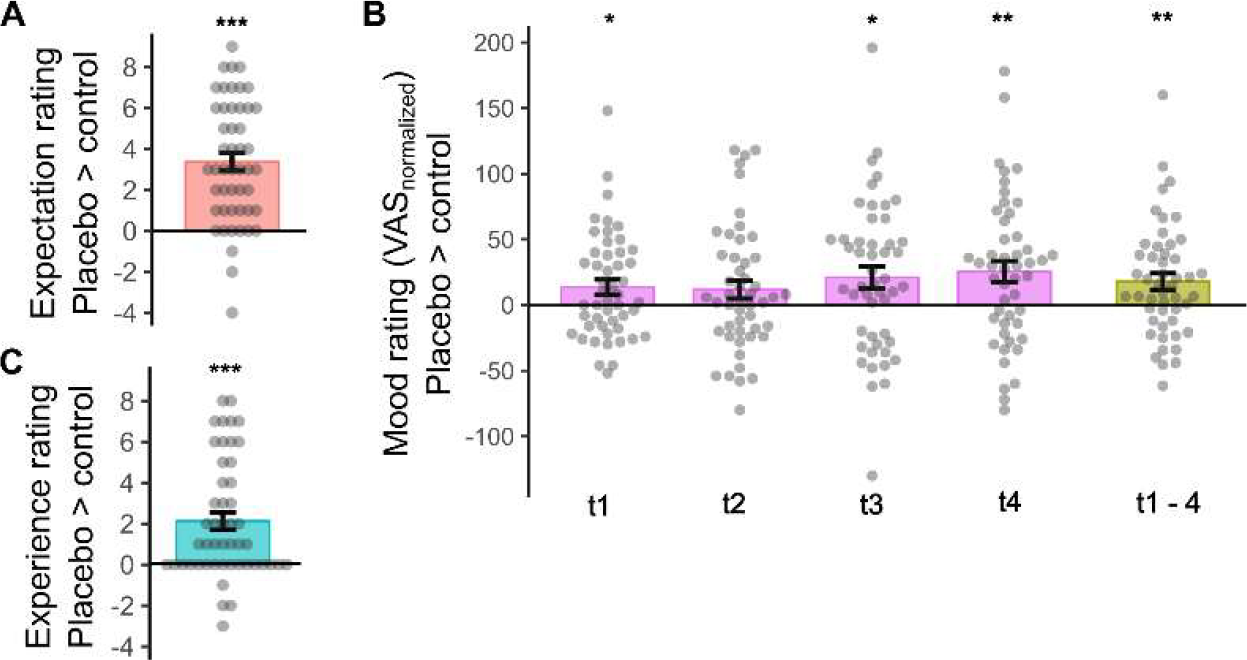
Placebo effects on expectation (A), mood (B), and experience ratings (C). Individual data with group means and s.e.m. Effects in baseline-normalized mood ratings are depicted for each time point and as an average across time points (t1 – 4). *p < .05, ** p < .005, *** p < .001.

Baseline mood state (VAS_t0_) did not differ between sessions (p = 0.12). rmANOVA on baseline corrected VAS values, considering ‘condition’ (control/placebo) and ‘time’ (t1-t4) demonstrated a strong condition effect (F_(1,48)_ = 8.61, p = .005, η^2^ = 0.15, Figure 2B), with higher mood ratings in placebo (mean difference of baseline corrected ratings VAS_t1-4_= 18.0 ± 6.1). There was no significant condition x time interaction (p > .26).

At the end of the sessions, participants reported a stronger experience of mood increase in the placebo session (T_(48)_ = 5.14, p < .001, d = 0.73, Figure 2C). Experience of mood enhancement at placebo was directly correlated with expectation ratings (r = .36, p = .012) as well as with positive changes in VAS mood ratings compared to control (r = .31, p = .029). There was no significant correlation between expectancy and VAS ratings (all p > .06).

### Placebo reduces distractibility by fearful faces in the high attention context

On both study days, participants showed high accuracy in task performance (errors: control: 3.0% ± 7.6%; placebo: 1.3% ± 2.0%). Replicating previous findings (36,37), attention was strongly manipulated by different cues, leading to significantly increased reaction time (RT) under the non-directing cue (NC) compared to the valid, directing cue (DC) condition in both the control (T_(48)_ = 12.09; p < .001, d = 1.73) and placebo sessions (T_(48)_ = 10.66; p < .001, d = 1.52), with no significant difference between sessions (F_(1,48)_ = 0.04; p > .85).

Eye-movement data of 27 participants could be analyzed (72% valid trials on average), showing that participants were well able to maintain central fixation across conditions (all p > .05; Supplementary Figure 2).

After-scan arousal ratings on the control day showed a significant effect of emotion (F_(3,46)_ = 37.45; p < .001, η^2^ = 0.71), caused by the following ranking of arousal ratings: happy ≈ fearful > sad > neutral. Ratings did not differ between sessions (F_(3,46)_ = 1.28; p > .29). As expected, post-scan valence ratings strongly matched pre-defined emotional expressions with an overlap of 94% on each of both sessions.

In an rmANOVA assessing baseline interference by emotional faces (neutral vs. fearful/happy/sad) under varying attentional conditions (high vs. low), a significant difference emerged only for fearful faces (F_(48,1)_ = 12.64, p < .001, η^2^ = 0.21), indicated by longer RTs. No significant effects were found for happy or sad faces, nor any emotion × attention interactions (all p > .16; Supplementary Figure 3).

To address the potential influence of intensity differences on this bias for fearful faces, we directly compared fearful to arousal-matched happy trials in another rmANOVA. Again, a significant main effect of emotion was found (F_(1,48)_ = 9.97, p = .003, η^2^ = .17), together with a significant emotion × attention interaction (F_(1,48)_ = 4.69; p = .035, η^2^ = .09). Slower responses in fearful versus happy trials, particularly in the high attention condition, drove these effects (T_(48)_ = 3.47, p < .001, d = .50; Figure 3). This bias for fearful faces in the high attention condition significantly decreased under placebo (T_(48)_ = 2.23, p = .031, d = 0.32), with no significant change in the low attention condition (p > 0.85). Accordingly, the abovementioned negativity bias for fearful versus neutral faces disappeared under placebo (p > .59), and still no bias was observed for happy or sad faces (all p > .82, see also Supplementary Figure 2). There were no placebo effects in control trials (scrambled faces across and within attentional conditions, all p > 0.81), indicating that placebo had no general effect on task performance.

**Figure 3.**
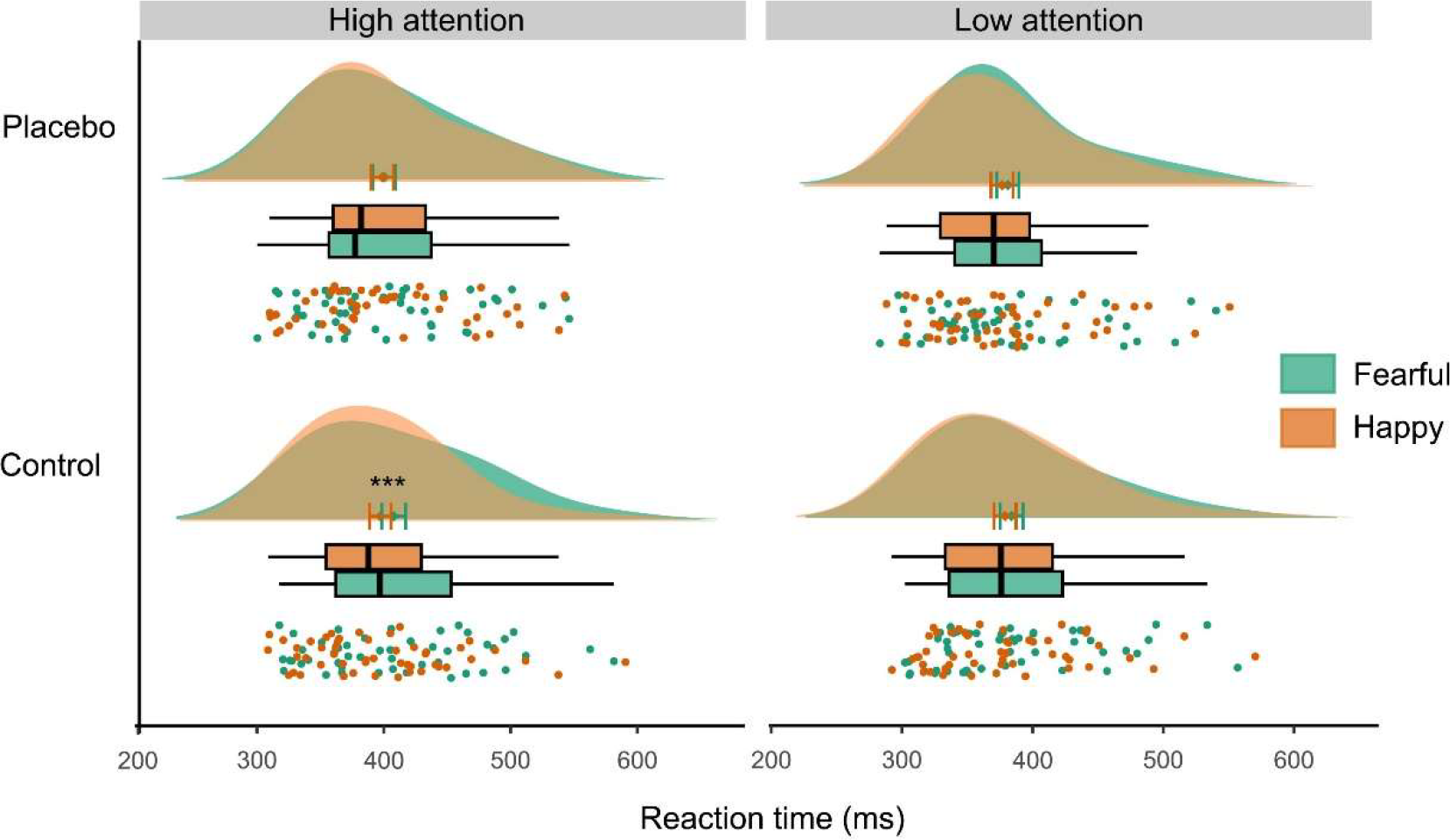
Behavioral results in the emotional interference paradigm. Distraction by fearful compared to happy faces at control (***p < .001) was significantly reduced at placebo (interaction: p < .04). Findings were restricted to the high attention to faces condition. Graph depicts individual data, boxplots, mean with s.e.m., and data distribution.

### Behavioral placebo effects are mediated by individual cognitive control ability

To investigate the potential modulation of placebo effects by participants’ expectation ratings and cognitive control ability, two ANCOVAs were conducted with session (control vs. placebo), emotion (fearful vs. happy), and attention to faces (high vs. low), along with covariates (expectation ratings / Singleton score). No significant interaction was found with subjective expectation ratings, but a significant relationship emerged between individual Singleton scores and the session by attention interaction (F_(1,47)_ = 10.09, p = .003, η^2^ = 0.18). Post-hoc correlations revealed that this effect was due to a specific correlation of cognitive control with placebo effects on highly attended fearful faces (control _fearful_nc_ > placebo _fearful_nc_ × Singleton: r = -.41, p = .004, Figure 4), indicating that individuals with higher cognitive control exhibited a more pronounced bias reduction.

**Figure 4.**
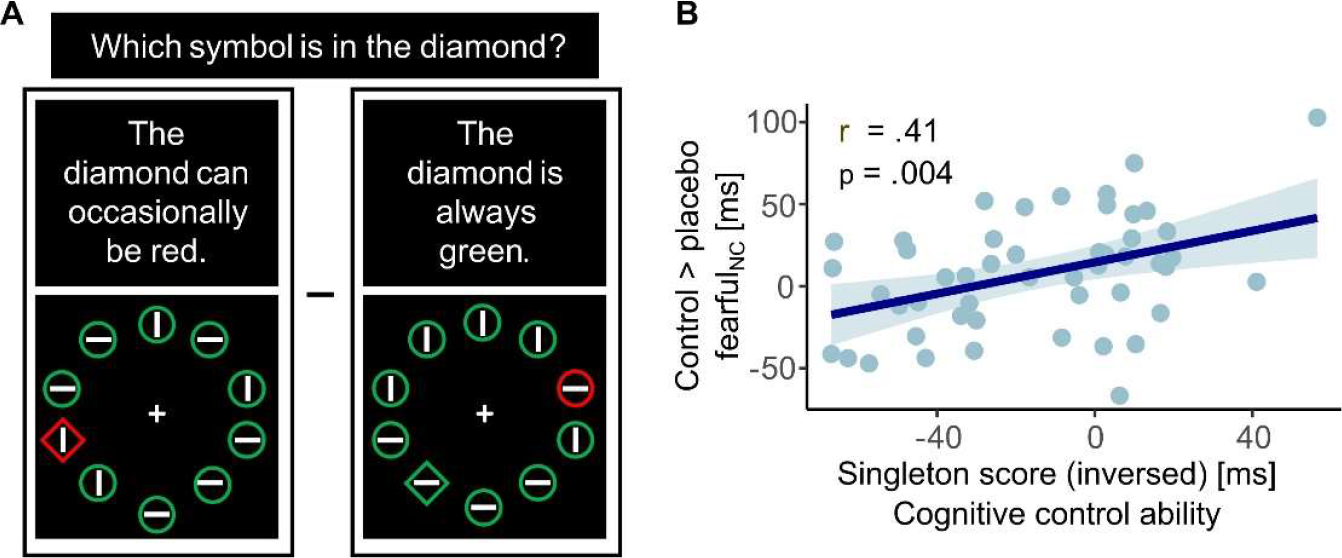
Correlation between behavioral findings and cognitive control ability. **(A)** Illustration of the two singleton-task conditions (for calculation of the singleton score, see Methods section) **(B)** Placebo effects on distractibility by fearful faces correlated with cognitive control ability.

### Neural correlates of fearful face distraction decrease at placebo

Initial confirmatory analyses validated the task’s attentional manipulation by demonstrating heightened activation in primary and higher order visual processing networks in response to non-directed cues compared to directed cues. Additionally, fMRI-data showed increased activity in a visual emotional network (14), involving the extrastriate visual cortex and bilateral amygdala, in response to face stimuli compared to scrambled pictures (Supplementary Figure 4). These patterns did not differ between sessions.

In our targeted analyses, we concentrated on fearful and happy face trials, mirroring the primary behavioral outcomes. We constructed an ANOVA for fearful versus happy faces, incorporating contrast images of high versus low attention to emotional faces from both sessions (control vs. placebo), resulting in a 2 × 2 × 2 factorial design. Consistent with behavioral findings, increased activity in a widespread network in response to fearful faces compared to happy faces was observed on the control day, likely indicating heightened distractibility by fearful faces. Regions involved cortical midline structures, such as the middle cingulate cortex and the bilateral precuneus, as well as superior frontal and parietal regions (Supplementary information, Table S1).

We then examined if there was a placebo-induced reduction in the response to highly attended fearful face stimuli (control > oxytocin: fearful_nc_ > happy_nc_). Results revealed significantly decreased activation under placebo in the anterior and middle cingulate cortex, left superior frontal gyrus, and precuneus, (Figure 5A). These effects were specifically observed in the high attention condition, with no activation differences in the low attention condition on the control day or in comparison to the placebo condition (see Table S1).

**Figure 5.**
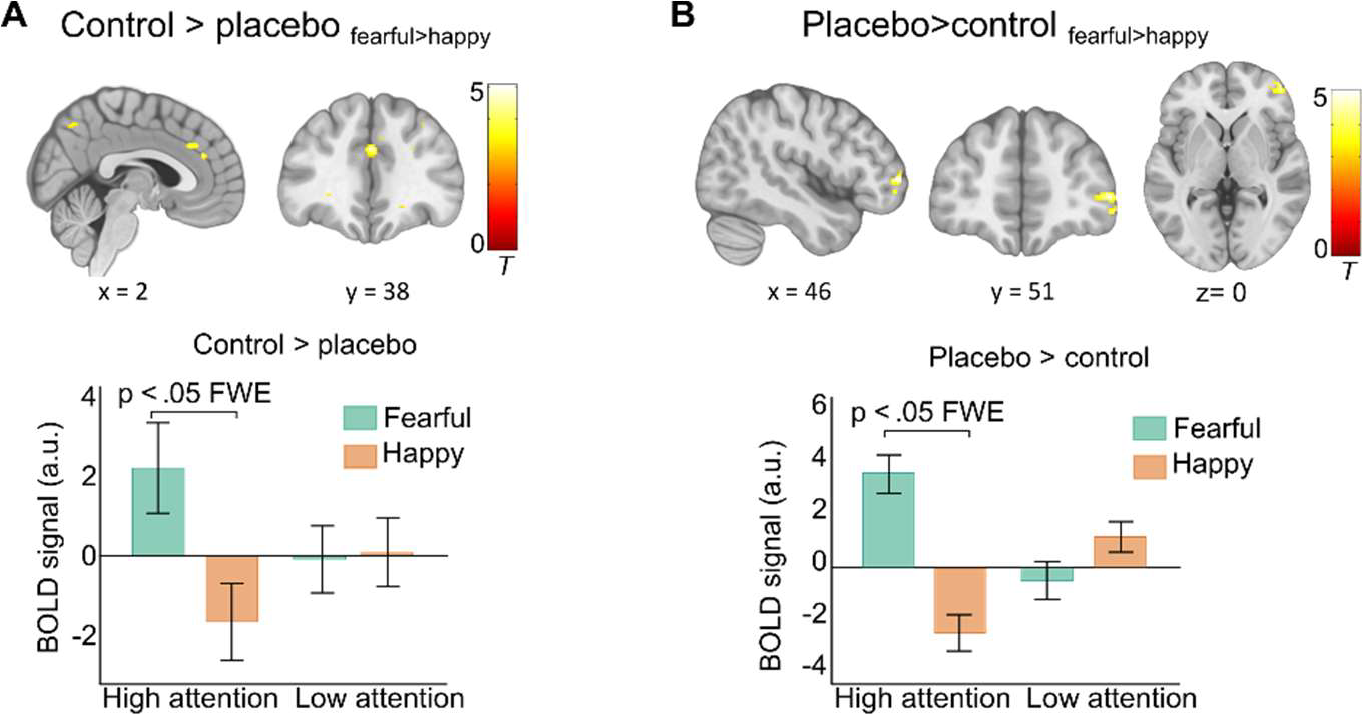
Neural placebo effects. **(A)** Placebo-induced signal decrease in response to highly attended fearful faces in the anterior and middle cingulate cortex, and precuneus. The bar graph shows group means and s.e.m of parameter estimates extracted from the middle cingulate cortex as examples of response patterns **(C)** Placebo-induced signal increase in the right middle frontal gyrus. The bar graph shows group means and s.e.m. of mean parameter estimates extracted from individual spheres within this activation cluster. All results are p < .05 FWE corrected. Activations are overlaid on a standardized MNI template (display threshold p < .001 uncorrected).

### Prefrontal modulation of the placebo effect

We next tested for potentially increased activation under placebo (placebo>control: fearful_nc_ > happy_nc_). Results revealed a significant cluster in the right anterior-lateral PFC, specifically in response to highly attended fearful faces (Figure 5B, Supplementary Table S1).

We employed psychophysiological interaction (PPI) analysis to examine if the functional coupling of this prefrontal region changed under placebo compared to control during the processing of highly attended fearful faces. Results indicated a stronger negative coupling between the lateral PFC and the ACC/vmPFC as well as the bilateral amygdala under placebo (Figure 6A, Supplementary Table S2).

**Figure 6.**
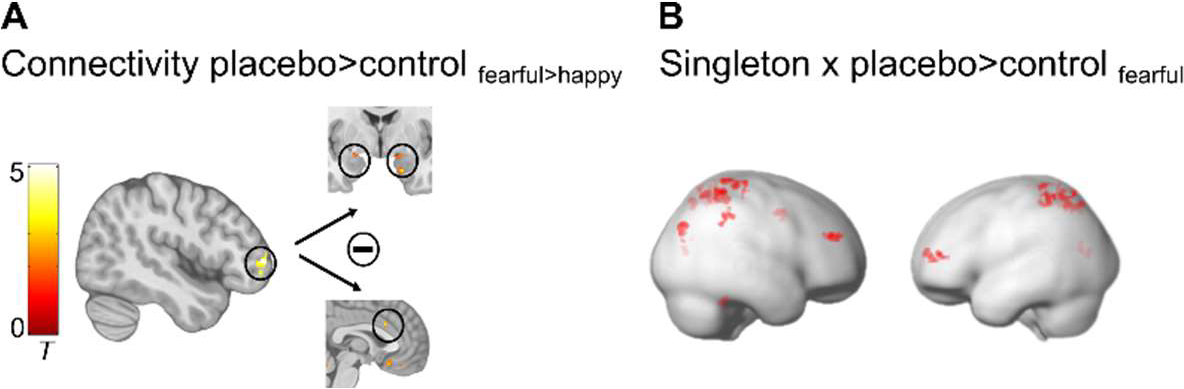
PPI and regression results. **(A)** Placebo-induced increased negative coupling of the right middle frontal gyrus with the bilateral amygdala and with the dorsal part of the ACC. **(B)** Cognitive control ability (Singleton) correlated positively with placebo-induced increased signals in a prefrontal-parietal network. All results are p < .05 FWE corrected. Activations are overlaid on standardized structural and rendered templates (display threshold p < .005 uncorrected)

### Placebo-induced prefrontal-parietal activity correlates with individual ability to control attention

Finally, to examine the neural correlates of the observed link between placebo effects and cognitive control capacity, we conducted a regression analysis including the contrast images fearful_placebo vs. control_ from the high attention condition and individual singleton scores as a covariate. Significant correlations emerged in the right dlPFC and the bilateral superior parietal lobe (SPL, Figure 6B and Supplementary Table S3). Thus, individuals with better attentional control showed a stronger placebo-related increase in prefrontal-parietal networks when being confronted with fearful face distractors.

## Discussion

The current findings reveal that placebo treatment, combined with positive instructions, improves subjective mood, and diminishes the processing of irrelevant aversive stimuli in a context with high attentional resources. This reduction in negativity processing is evident through decreased activation within emotional interference networks, accompanied by heightened involvement of the lateral PFC and its effective down-regulation of limbic appraisal networks. Importantly, participants’ overall cognitive control capacity directly predicted both behavioral and prefrontal-parietal placebo effects. These results highlight the significance of resource-dependent top-down regulation in verbally instructed placebo effects on affective processing, with implications for the potential and constraints of treatment expectations, particularly within clinical contexts.

The placebo-related decrease in aversive stimulus perception aligns with prior findings on placebo analgesia and affective expectation manipulation. Many studies used a combination of verbal instructions and conditioning to maximize expectation effects (14,44,45). Our study demonstrates that such a decrease can also be achieved through verbal instruction alone. Neurophysiologically, diminished distraction by fearful face distractors was paralleled by an active suppression of neural networks processing emotional conflicts (46,47), including the middle cingulate cortex, the superior prefrontal cortex, and the precuneus. Reduced BOLD signals in these regions indicate decreased interference due to reduced attentional distraction under placebo.

Verbally instructed placebo effects have been linked to changes in expectations and appraisal processes, primarily mediated by prefrontal brain networks modulating subcortical, limbic systems (26,48,49). Our data support these notions, revealing reduced processing of fearful faces associated with enhanced activity in the right lateral PFC. Connectivity analyses further indicated an increased negative coupling of this region with the amygdala and the vmPFC during placebo treatment, suggesting enhanced inhibition of affective valuation processes by higher-order cognitive control. The lateral PFC, crucial for cognitively demanding explicit strategies and cognitive flexibility (50), has been associated with the top-down biasing of affective key nodes according to the placebo-instructed state (26). This engagement may be modulated through the vmPFC, integrating information from the lateral PFC about current goals with subcortical input about stimulus relevance from the amygdala (25,26,51,52). Thus, our data suggest that increased lateral PFC activity signals instructed beliefs about the positive effects of oxytocin and biases appraisal processes in downstream networks accordingly through the inhibition of value signals in response to aversive stimuli.

We focused on the role of cognitive resources in verbally instructed placebo effects within the affective system. Despite striking evidence for an involvement of prefrontal-limbic top-down regulation in placebo effects across systems (25,26,53,54), results on direct modulation through cognitive processes are heterogeneous. For instance, previous data suggested that placebo analgesia does not rely on an active redirection of attention and is unaffected by cognitive distraction (30). Conversely, reduced executive functioning and weaker prefrontal resting-state EEG connectivity have been associated with disrupted placebo responses in Alzheimer’s disease patients (29) and cognitive reappraisal ability mediated via the dorsolateral PFC has been positively related to placebo analgesia in young adults (31). To our knowledge, no study so far has investigated the role of cognitive resources in affective placebo effects.

We examined the impact of cognitive resources in two ways: first, as context-dependent state variable and, second, as a context-independent general capacity. Our results showed that high resources in both dimensions predicted stronger placebo effects. Replicating previous findings (36,37), our experimental paradigm effectively modulated attentional resources for processing emotional distractors, evident in prolonged reaction times and amplified neural face processing in the high compared to the low attention condition. Placebo effects selectively manifested in trials with non- directing cues, where distraction might have been strongest during the control session, but more resources were available for voluntary attentional allocation during the placebo session. This aligns with prior research using this paradigm, showing a positivity bias in older adults confined to the high attention condition (37). In contrast to a previous study on positive expectation effects in emotion detection (19) we did not observe a positivity bias in terms of increased distraction by happy faces under placebo. This could be attributed to divergent task instructions, specifically the distinction between emotional detection and interference. Also unlike our earlier work (19), we observed that expectation ratings in this study were only correlated with perceived mood improvement and not with neurobehavioral placebo effects. This discrepancy might arise from individual variations between objective and subjective measures of placebo effects, as well as potential limitations in the validity of our expectation assessment. Although our manipulation significantly affected expectation ratings, they were assessed only once after nasal spray application to prevent participants from becoming suspicious through repeated ratings. While this approach aimed to maintain experimental integrity, it may have limited overall validity when relying on a single, moderately scaled rating for the entire experiment.

The selective allocation of attentional resources in the service of prioritized emotional goals in a trial-by-trial manner (41,55) likely depends on individuals’ general capacity to flexibly control their attention. Here we investigated the basic principle of this assumption by assessing individuals’ general cognitive ability to exert attentional top-down control over salient distraction with a Singleton visual search task (56,57). This task evaluates the specific function of spontaneously focusing on target information over salient distraction, and has previously been related to peoples’ ability to voluntarily control their attention in an emotional context (41). Both, behavioral and neural placebo effects were directly correlated with participants’ performance in this task. At the neural level, control ability was associated with a specifically activated network of visuo-spatial orienting and top-down control (58,59), comprising the right dorsolateral PFC and the bilateral superior parietal lobe (SPL).

Our findings may have important implications for the understanding of expectation effects in the treatment of mood disorders with antidepressants, both in the context of active placebo treatment and as beneficial “add-on” to effective treatments (1,60,61). Although prioritizing negative information is a frequent finding in healthy individuals (62–65), a pathologically enhanced focus on negative information is a key feature in clinical depression and is crucial for the persistence and prognosis of depressive symptoms (66–68). It therefore seems intriguing that through verbal instruction alone, attentional preferences for emotional stimuli can be modified together with mood state improvements and positive treatment experience, with the latter being an important prior for future treatment success. Whether this also applies to clinical samples and how stable such effects are over time must be addressed in future studies. Patients with major depressive disorder often demonstrate deficits in prefrontally mediated cognitive control (32–35). Our findings suggest that patients with those deficits may benefit less from verbally instructed expectations. Their expectations might be boosted by conditioning, learning and positive experiences by which more implicit, cognitively less demanding regulation could be achieved (69,70). First studies have started to develop such procedures (4,15,23). Together with the consideration of individual cognitive control ability this may pave the way for an individually tailored intervention in the treatment of affective disorders.

## End notes

## Author contributions

Concept and design: S.B., Acquisition of data: J.R., A.M., Analysis or interpretation of data: J.R., M.G., C.B., W.R., S.B., Drafting of the manuscript: S.B., A.M., Critical revision of the manuscript: All authors.

## Acknowledgements

We thank F. Thams, J. Baker and E. Shin for their assistance with data collection. The work was funded by grant CRC 289 Treatment Expectation, Project-ID 422744262, from the Deutsche Forschungsgemeinschaft (DFG, German Research Foundation)

## Data availability

Behavioral and imaging data that support the findings of this study have been deposited online under the following link: https://doi.org/10.6084/m9.figshare.24279538.v1

