## Supplementary information for "Placebo treatment entails resource-dependent downregulation of negative inputs"

### Analysis of eye movement data

Eye-tracking data were first parsed into rapid eye movements exceeding a velocity of 30°/s or an acceleration of 8000°/s^2^ (saccades) and relatively stable gaze periods between the saccades (fixations). Subsequently, we subtracted the last 300 ms before stimulus onset from fixation coordinates to relate individual gaze positions to baseline values where the centrally presented cross should be fixated. To ensure that participants adhered to this instruction, we used an iterative outlier detection algorithm on x- and y-coordinates of baseline values (2). Specifically, the highest and lowest values of baseline data were temporally removed from the data set, and it was tested whether they deviated more than three SDs from the mean of the remaining distribution. If one or both values met this criterion, they were marked as outliers and excluded from the data set, if not, they were returned. This was iteratively repeated until no further x- and y-values had to be removed. Trials with blinks during the baseline period were also removed.

After trial selection and drift correction, we identified all fixations that fell on the left (x < 0) and right side of the screen (x > 0) and calculated the total dwell time on both sides during the entire cueing period before the target appeared. From these values, we derived the proportion of dwell time on the screen side that was cued (for directional cues) or on the right side of the screen (for neutral cues) by dividing dwell times by the total duration of all valid fixations in this period. These values were averaged for all trials of the same condition and compared using a 2 (day: placebo vs. control) × 5 (face: fearful, happy, sad, neutral, scrambled) × 2 (cue: non-directed vs. directed) ANOVA. For illustrative purposes, we also calculated trajectories of horizontal gaze displacements from fixation coordinates in bins of 10 ms for the whole cueing period.

### Singleton task

On each trial, nine circles (distractors) and one diamond (target shape) were presented equally spaced around the fixation point. Participants had to indicate as fast and accurately as possible whether the target shape (diamond) contained a horizontal or a vertical line, using dedicated keys on a standard computer keyboard. There were two conditions indicated by different instructions. In the baseline condition (BC), participants were informed that the target-diamond is always green. Specifically, 50% of all trials contained no particularly salient item (i.e., no singleton, all items green, BC_all___green_), while in the other 50%, one distractor (circle) was printed in red (singleton distractor, BC_singleton_distractor_). Thus, in the baseline condition participants could completely concentrate on one color. In the Singleton-detection condition (SC), participants were instructed that the color singleton could also be the target. In fact, in ~8% of trials the target-shape was printed in red (singleton target), in addition to trials with no singleton (SC_all_green_) and singleton distractors (SC_singleton_distractor_). That is, now the singleton color could not be easily ignored (see Figure 4A). The experiment consisted of 8 blocks, 4 for each condition, presented alternately. Each block contained 48 trials, with instructions displayed at the beginning and midway through each block. Each trial began with a fixation period of 1.1 to 1.25 seconds, followed by stimulus presentation for up to 3 seconds, and a random inter-trial interval ranging from 0.2 to 0.8 seconds. Feedback was provided for either a failure to respond within the 3-second window or an incorrect response, with messages displayed as “too slow” or “error”, respectively. All trials in which volunteers responded inaccurately or faster than 200 ms were excluded. Trials with a singleton target and the subsequent trial were also excluded due to specific response patterns induced by such trials. The individual singleton score was then calculated using reaction times by the following equation:

Singleton score = [*SC*_singleton_distractor_ – *SC*_all_green_] – [*BC*_singleton_distractor_ – *BC*_all_green_]

Thus, a higher singleton score implied less flexible attentional control.

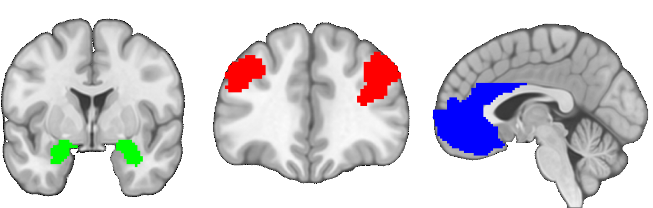

**Supplementary Figure 1. Regions of interest (ROI).** Predefined ROIs in the amygdala (green), dlPFC (red), and the vmPFC/ACC (blue) overlaid on the MNI structural template. ROIs were created based on functional clusters (p < 0.05 FDR corrected) from meta-analyses on neurosynth.org using “DLPFC” (489 studies) and “VMPFC” (199 studies; August 2023) as search terms. Due to reported overlapping placebo results in vmPFC and ACC, the vmPFC mask was combined with the ACC mask from the AAL3 atlas (1). Bilateral amygdala masks were also derived from AAL3.

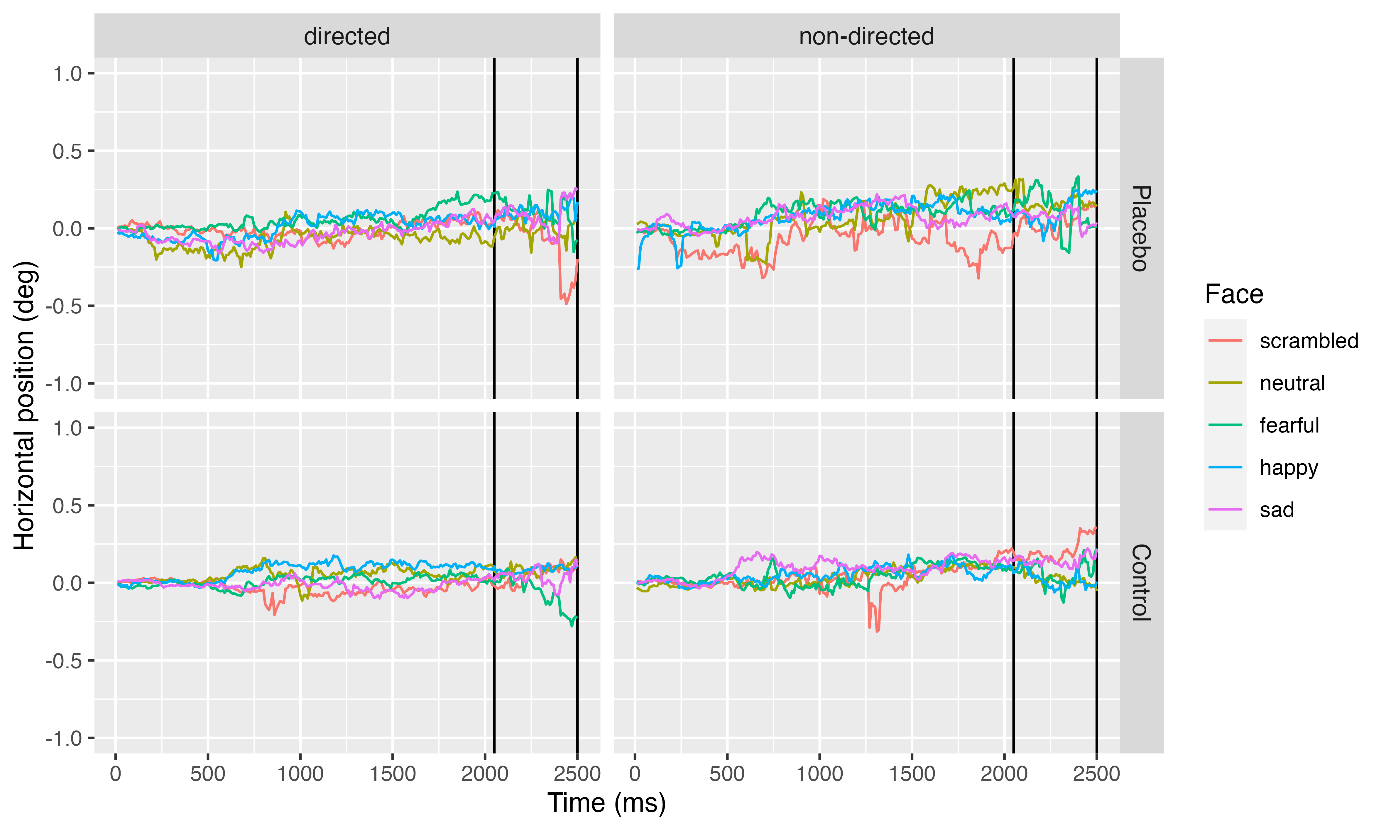

**Supplementary Figure 2. Horizontal gaze trajectories as a function of face stimulus and cue for the placebo and the control day.** Data are depicted in degrees of visual angle with positive values indicating gaze shifts into the direction of the cue (directed cues) or towards the right side of the screen (neutral cues). Vertical lines denote the time interval during which the cueing period ended. Gaze shifts did not differ significantly between conditions and were very small considering that the face stimulus spanned approximately 5° horizontally.

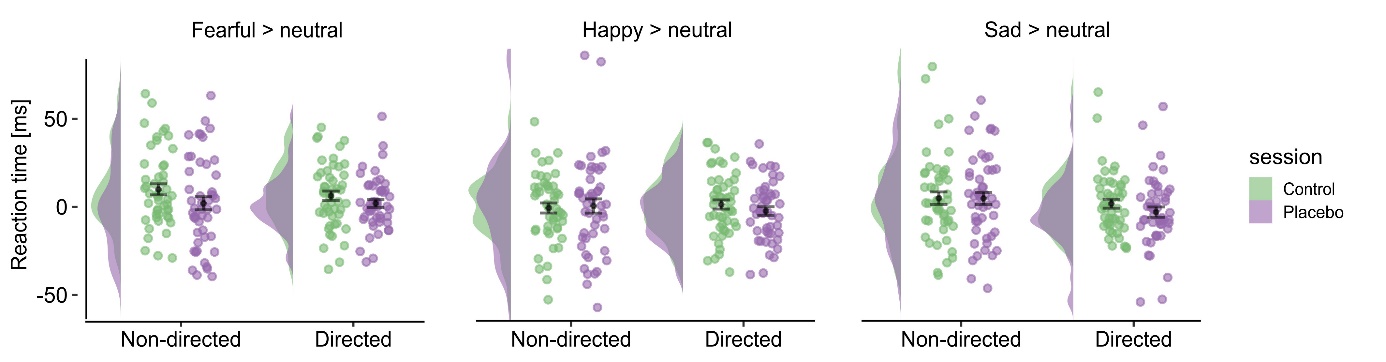

**Supplementary Figure 3. Reaction times for each emotional expression compared to neutral faces for both cues and sessions.** All plots include means, SEM, individual data and density plots.

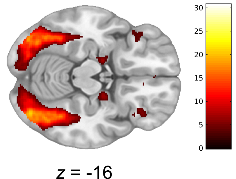

**Supplementary Figure 4. Visual emotional network including the extrastriate cortex and the bilateral amygdala in response to face stimuli as compared to scrambled pictures**

|  |  | MNI (peak) | | |  |  |
| --- | --- | --- | --- | --- | --- | --- |
| Brain region | Side | x | y | z | Cluster size | Z |
| **Main effect emotion, control session** |  |  |  |  |  |  |
| *Fearful > happy, non‐directed cue* |  |  |  |  |  |  |
| Superior frontal medial gyrus / middle cingulate cortex | R | 3 | 29 | 34 | 106 | 4.42 |
| Middle temporal gyrus | L | ‐64 | ‐25 | ‐6 | 155 | 5.84 |
| Inferior parietal gyrus | L | ‐36 | ‐50 | 47 | 94 | 4.24 |
| Precuneus | R | 10 | ‐67 | 42 | 203 | 5.20 |
|  | L | ‐12 | ‐67 | 46 | 76 | 5.10 |
| *Fearful > happy, directed cue* |  |  |  |  |  |  |
| n.s. |  |  |  |  |  |  |
| *Happy > fearful, non‐directed cue* |  |  |  |  |  |  |
| Putamen | L | ‐27 | ‐1 | 5 | 75 | 4.18 |
| Angular Gyrus | R | 51 | ‐62 | 22 | 76 | 4.55 |
| Calcarine cortex | L | ‐12 | ‐91 | ‐4 | 166 | 5.04 |
| Cuneus | R | 20 | ‐94 | 10 | 226 | 5.61 |
| *Happy > fearful, directed cue* |  |  |  |  |  |  |
| n.s |  |  |  |  |  |  |
| **Placebo eﬀects** |  |  |  |  |  |  |
| *Control > Placebo: fearful > happy, non‐directed cue* |  |  |  |  |  |  |
| Superior frontal gyrus | L | ‐28 | 42 | 23 | 89 | 4.07 |
| Anterior cingulate cortex |  | 0 | 38 | 26 | 55 | 4.79* |
| Middle cingulate cortex | R | 8 | 23 | 30 | 132 | 4.61 |
| Precuneus | R | 10 | ‐67 | 44 | 128 | 4.98 |
| *Control > Placebo: fearful > happy, directed cue* |  |  |  |  |  |  |
| n.s. |  |  |  |  |  |  |
| *Placebo > Control: fearful > happy, non‐directed cue* |  |  |  |  |  |  |
| Middle frontal gyrus | R | 46 | 52 | 2 | 150 | 4.95 |
| *Placebo > Control: fearful > happy, directed cue* |  |  |  |  |  |  |
| n.s. |  |  |  |  |  |  |
| **Supplementary Table 1 \| Peak coordinates and statistics for fMRI activations**  Montreal Neurological Institute (MNI) coordinates and the respective z-values are reported for peak voxels and local maxima within each cluster. All p < .05 FWE corrected. * Significant results based on small volume corrections. L: left, R: right, n.s.: not significant | | | | | | |

|  |  | MNI (peak) | | |  |  |
| --- | --- | --- | --- | --- | --- | --- |
| Brain region | Side | x | y | z | Cluster size | Z |
| **Control > placebo, feaful > happy, non‐directed** | |  |  |  |  |  |
| Anterior cingulate cortex | L | -2 | 18 | 29 | 28 | 4.35* |
| Amygdala | L | ‐24 | ‐4 | -14 | 12 | 3.58* |
|  | R | 27 | 6 | -19 | 9 | 3.61* |
| **Placebo > control for fearful > happy, non‐directed** |  |  |  |  |  |  |
| n.s. |  |  |  |  |  |  |

**Supplementary Table 2 | PPI results** Montreal Neurological Institute (MNI) coordinates, cluster size and respective z-values are detailed for the gPPI analysis, utilizing the right middle frontal gyrus (46,52,2) as a seed region. All p < .05 FWE corrected. * Significant results based on small volume corrections. L: left, R: right, n.s.: not significant.

|  |  | MNI (peak) | | |  |  |
| --- | --- | --- | --- | --- | --- | --- |
| Brain region | Side | x | y | z | Cluster size | Z |
| **Placebo > control, fearful, non‐directed** × **singleton** | |  |  |  |  |  |
| Middle frontal gyrus | R | 40 | 44 | 23 | 52 | 4.28* |
| Superior parietal gyrus | L | ‐26 | ‐58 | 53 | 66 | 4.27 |
|  | R | 36 | ‐50 | 60 | 64 | 3.98 |
| **Supplementary Table 3 \|Simple linear regression analysis with singleton-score as covariate**  Montreal Neurological Institute (MNI) coordinates, cluster size, and respective Z-values are reported for peak voxels and local maxima within each cluster. All p < .05 FWE corrected. * Significant results based on small volume corrections. L: left, R: right, n.s.: not significant. | | | | | | |
